# Time-Dependent Facilitation of Homologous Actions

**DOI:** 10.1101/2025.10.15.682693

**Authors:** Raphael Hamel, Felix-Antoine Savoie, David Punt, Ned Jenkinson, Mark R. Hinder

## Abstract

Unimanual actions can interfere with or facilitate similar actions performed with the opposite hand, especially when in close temporal proximity. Across three sequential button-press experiments, we tested how effector homology – anatomical similarity between fingers – and temporal delays between actions shape these effects. Specifically, we examined whether a priming action altered the reaction time (RT) of a subsequent action. Compared with unimanual RTs, we indexed slowing of the second action’s RT as interference, and quickening as facilitation.

Priming with homologous actions (e.g., index finger-index finger) produced interference at short intervals (≤200ms) but transitioned to facilitation at longer intervals (≥400ms). Priming with non-homologous actions (e.g., little finger-index finger) also produced interference at short intervals, but never resulted in facilitation. Critically, these patterns emerged whether the priming actions were performed with the opposite or the same hand, indicating that interference and facilitation do not depend on interhemispheric dynamics.

Our results reveal a previously undocumented time-dependent shift from interference to facilitation that is specific to homologous actions, challenging models that explain the interference between parallel actions solely by competitive interhemispheric dynamics or central bottleneck processes. We propose that facilitation and interference flexibly coexist, and are shaped by effector homology and timing. These findings extend current models of bimanual coordination and highlight new opportunities for enhancing motor performance and neurorehabilitation.

**GRAPHICAL ABSTRACT:** A left-hand index finger press primed a subsequent right-hand index (homologous) or right-hand little finger (non-homologous) response at variable delays. For homologous actions (index–index), reaction times (RTs) quickened relative to unimanual single-press baseline for delays ≥400ms, whereas for non-homologous actions (index–little), RTs slowed and gradually returned to baseline by ~600ms. This dissociation indicates that effector homology and timing dictate whether a preceding action facilitates or interferes with the execution of a subsequent one.

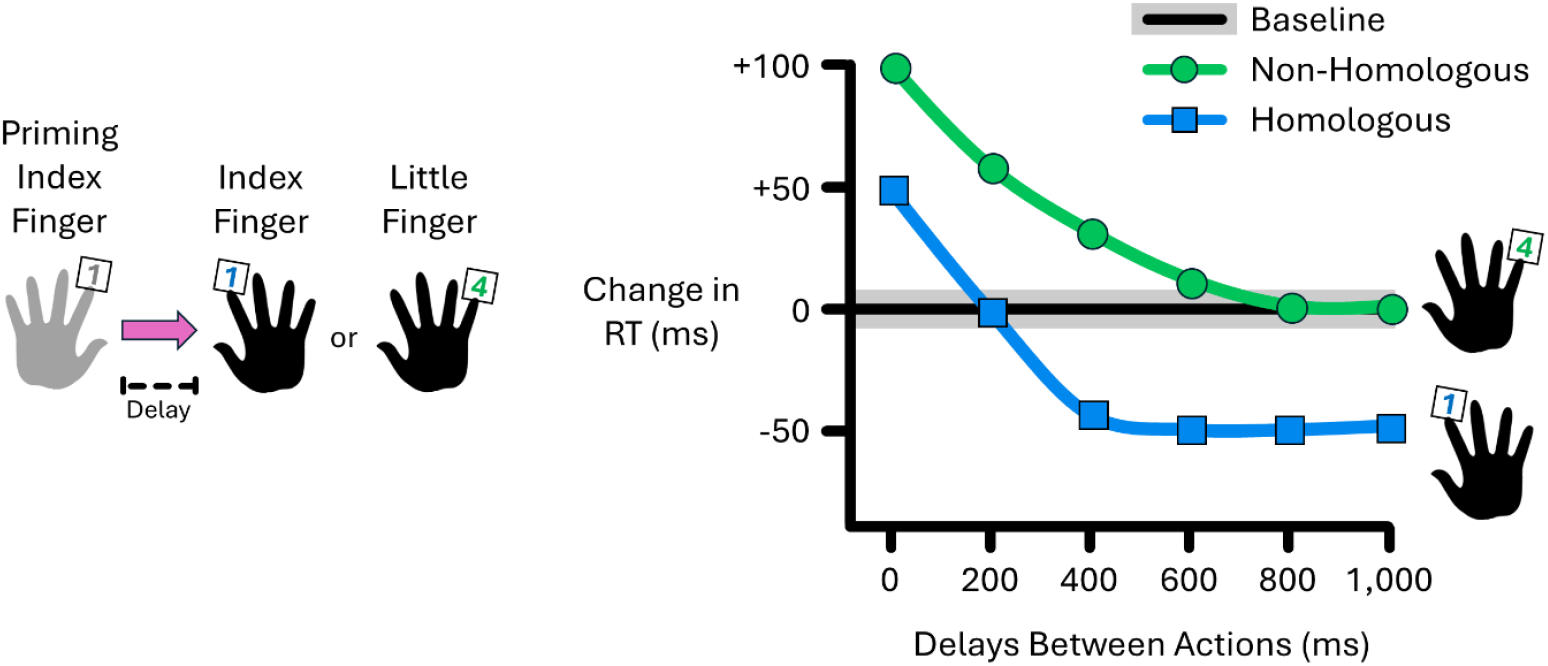

## INTRODUCTION

We often use both hands to perform sequential actions, such as tying shoelaces or typing on a keyboard. But how does the brain generate such bimanual actions? One influential view holds that a unimanual action triggers interhemispheric inhibition, suppressing excitability in the opposite homologous effector via transcallosal projections [1]. This mechanism may help prevent bimanual mirror movements and competing responses [1]. However, it also hinders performance when the contralateral effector performs discrete actions simultaneously [2,3] or with a delay [4,5]. According to this evidence, the control of two concomitant unimanual actions is largely governed by interhemispheric competition.

An alternative view proposes that unimanual actions trigger interhemispheric facilitation – rather than inhibition – in the opposite homologous effector [6]. In support, discrete unimanual actions increase excitability [7–9] and reduce inhibition in the motor cortex controlling the contralateral homologous effector [10], suggesting that bimanual actions recruit facilitatory interhemispheric mechanisms. Such facilitatory processes have been proposed to enhance motor output – particularly force production – in bimanual tasks [11,12]. Yet, whether such facilitation plays a functional role in bimanual action control beyond enhancing force production remains unknown. It is also unclear whether interference and facilitation can coexist, and whether inhibition can transition to facilitation – or vice-versa – depending on task demands.

Here, we propose that bimanual actions can trigger interference or facilitation depending on two key factors: effector homology and temporal delays. Homologous actions are supported by homotopic transcallosal projections [13,14] and intra-hemispheric recurrent loops [15], which may expedite action preparation and/or execution by facilitating synchronous neural activation of homologous effectors. In contrast, heterotopic projections between non-homologous effectors may be less direct and efficient [14], possibly incurring additional neural computations that increase susceptibility to interference. Based on this, one hypothesis is that homologous actions produce minimal interference and may even enable facilitation. Additionally, electroencephalography studies showed that sensorimotor brain activity – inhibitory-dominant beta-band rhythms – remains suppressed for a few seconds after completing a single action [16,17] and until the second action of a sequence has been performed [18]. Our second hypothesis directly builds on this evidence: a first action may temporarily disinhibit the motor system, which could time-dependently modulate the preparation of a subsequent action. Thus, we expected the pattern of interference and facilitation to depend on the timing of the two actions.

## METHODS

### Participants

Twenty neurologically healthy adults participated in each of our three experiments (total n = 60; 41 females; 57 right-handers; 27.5 ± 1.0 years old, Mean ± SEM). One participant took part in all three experiments, while seven others participated in two of the three. Each experiment was powered to detect a minimum within-subject effect size of Cohen’s *dz* = 0.66 (α = 0.05, two-tailed, 80% power; paired *t*-test). While this power estimate is based on a simplified model, each participant completed over 500 trials within each experiment, providing high data density for detecting reliable within-subject effects using trial-level generalised linear mixed models.

All participants provided informed consent prior to participation. The study was approved by the University of Tasmania Human Research Ethics Committee (project #31750) and conducted in accordance with the Declaration of Helsinki. Participants received either two hours of research credit or AUD $20 financial compensation for their time.

### Apparatus

All tasks were programmed using PsychToolbox-3 in MATLAB R2024a (MathWorks). Button presses were recorded using a Scorpion K636 membrane gaming keyboard (1ms response time). Visual stimuli were presented on a ViewSonic XG monitor with a 240Hz refresh rate. On the keyboard, participants responded using the left index finger on the “S” key (labelled “1”), the right index finger on the “G” key (also labelled “1”), and the right little finger on the “L” key (labelled “4”). Fingers remained in position over the assigned keys throughout.

### Experiment 1

Figure 1. provides an overview of a typical trial timeline (**Panel A**) and effector combinations across experiments (**Panel B**).

**Figure 1.**
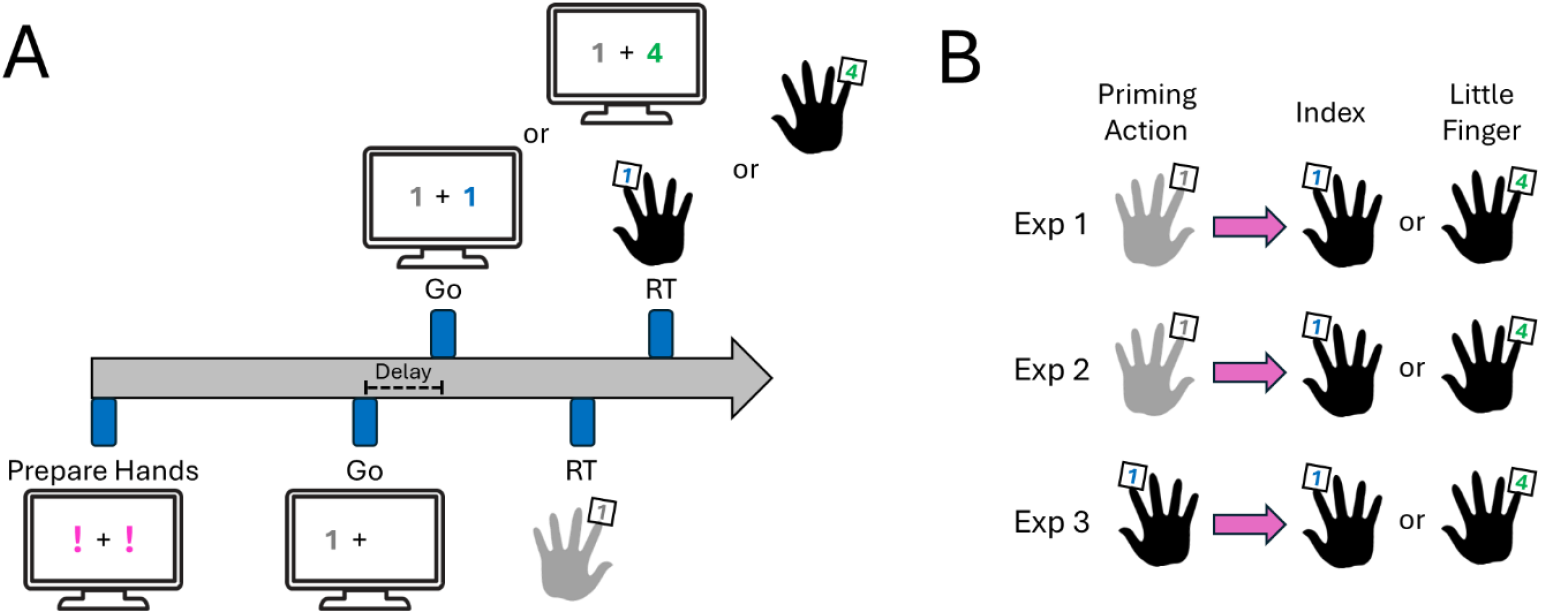
A first press primed a homologous right index finger or non-homologous right little finger press at variable delays: 0, 200, 400, 600, 800, 1,000ms. **(A)** The delays separated the Go visual stimulus of each action. **(B)** Priming presses consisted of left-(Exps 1 and 2) or right-handed (Exp 3) index finger presses. In Exp 2 **(B)**, simultaneous presses (0ms delay) were removed. In Exp 3 **(B)**, simultaneous homologous presses could not be performed due to same-finger use.

In Exp 1, we tested whether interference and facilitation effects depend on effector homology and temporal delays between two actions performed with separate hands.

Participants performed fast and accurate button presses in response to visual stimuli. Each trial began with a fixation cross and one or two magenta exclamation marks (“!”). A single “!” prompted a unimanual press, whereas two “!” indicated bimanual presses. When two “!” appeared, participants were never informed of the delay between the two presses. As shown in **Figure 1A**, the “!” spatial location served as cues for which hand/finger to prepare. Left-sided “!” invariably prompted simple RT presses from the left index whereas right-sided “!” prompted choice RT presses from the right index or right little finger. After 1,000ms ± 250ms (random jitter), the “!” turned into numerical Go stimuli: “1” and “4” digits prompted index and little finger presses, respectively.

Homologous presses involved the left and right index fingers, while non-homologous presses involved the left index and right little fingers (**Figure 1B; top**). To manipulate temporal delays, the numerical Go stimuli would either appear simultaneously (0ms delay) or sequentially at delays of 200, 400, 600, 800, or 1,000ms. On sequential trials, participants were informed that left index presses would always prime presses from right index or little finger.

After each response, accuracy (“Great!” in green when successful; “Oops” in red when failed) and reaction time (RT) feedback (“RT: 347ms”) were provided. Trials were categorised as failed if participants omitted a response, pressed the wrong key, or responded prematurely (<150ms). For simultaneous presses (0ms delay), trials were also marked as failed if the inter-press interval exceeded 25ms.

### Experiment 2

Exp 1 revealed that homologous presses quickened RTs relative to baseline values when delays were ≥400ms (see Results). Hereafter, we refer to this quickening as RT facilitation.

Exp 2 tested whether this facilitation effect depended on overt interhemispheric temporal coupling. One possibility is that this result was confounded by the presence of simultaneous (0ms) presses. Specifically, even though conditions were fully pseudorandomised across trials, the presence of simultaneous presses may have implicitly encouraged participants to adopt a temporally-coupled response mode across all trials [19,20]. Thus, the RT facilitation we observed may not solely reflect effector homology, but also a by-product of task demands emphasising temporal coupling between hemispheres.

To disentangle these possibilities, Exp 2 was identical to Exp 1, except that simultaneous press conditions were removed. Thus, only the five sequential delays remained: 200, 400, 600, 800, and 1,000ms. If facilitation persisted despite the absence of simultaneous presses, it would suggest that interhemispheric temporal coupling is not necessary for the effect. In that case, facilitation may instead reflect a more general property of motor system organisation, such as structural or representational similarity between homologous effectors [13,14]. Homologous and non-homologous finger combinations remained unchanged from Exp 1 (**Figure 1B, middle**).

### Experiment 3

Exp 3 was structurally identical to Exp 1, except that priming *left* index finger presses were replaced with *right* index finger presses. This allowed us to test whether interference and facilitation can emerge within a single hemisphere, by restricting all responses to the right hand. Left-sided “!” cues now prompted a simple RT press with the right index finger, while right-sided “!” continued to cue a choice RT press with either the right index or right little finger. Homologous presses thus involved two right index finger presses, whereas non-homologous presses paired the right index and right little fingers (**Figure 1B, bottom**).

The same set of temporal delays used in Exp 1 was applied, except that the 0ms delay condition was removed from homologous presses. This was because executing two simultaneous presses using the same finger is not physically possible.

### Data Processing and Analysis

Unimanual presses performed without preceding primes served as a baseline. Separate baselines were computed for each experiment and for each effector. RTs from the second actions – when primed by a first press – were then compared against these baselines to assess whether priming altered performance. RTs slower than baseline were interpreted as interference, whereas RTs faster than baseline were taken as facilitation.

RTs were calculated on a trial-by-trial basis as the time (ms) between the onset of the Go stimulus and the corresponding finger press. For trials involving two presses, RTs were computed separately for each response. Each participant completed a total of 42 trials per temporal delay and effector, which were equally distributed across three blocks. Within each block, trials were pseudorandomised so that conditions never repeated on adjacent trials. Across all experiments, 10.6% of all trials were failed and excluded from analyses.

Trial-level RTs were analysed without prior aggregation using generalised linear mixed models, which included a gamma distribution and log-link function to account for RT positive skewness [21]. RTs from the second homologous and non-homologous presses were analysed in separate models because they involved different responding effectors (right index finger vs right little finger) and, therefore, distinct unimanual baselines. Including homology as a factor would risk confounding baseline effector differences with homology effects. Our interest was not in directly contrasting homologous with non-homologous actions, but in testing whether each condition diverged from its respective unimanual baseline.

The fixed effect was Delays (Unimanual, 0ms, 200ms, 400ms, 600ms, 800ms, 1,000ms), with the Unimanual condition serving as the baseline for each delay. Random intercepts (participants) and random slopes (delays) were included in each model [22]. All generalised mixed models converged. Statistical significance was set at *p* < 0.05 throughout, with Benjamini–Hochberg corrections [23] applied to control for multiple comparisons. All analyses were conducted in JAMOVI (v2.4.14) using the GAMLj module [24]. Figures display model-derived estimated marginal means ± standard error (SE).

### Data availability

All data underlying the findings of this study are openly available. The datasets for the main and supplementary analyses are archived as files at https://osf.io/z8t5m/.

## RESULTS

### Exp 1: How did interference and facilitation emerge during bimanual actions?

**Figure 2A** shows that RTs during homologous actions varied with the delay between left and right index finger presses (χ^2^ = 177, df = 6, *p* < 0.001). The right index finger showed RT slowing of ~50ms during simultaneous presses (*p* < 0.001), which returned to baseline at 200ms (*p* = 0.199), and became ~50ms faster than baseline for delays ≥400ms (all *p* < 0.001). This shows that the timing between two homologous actions determines whether interference or facilitation occurs.

**Figure 2.**
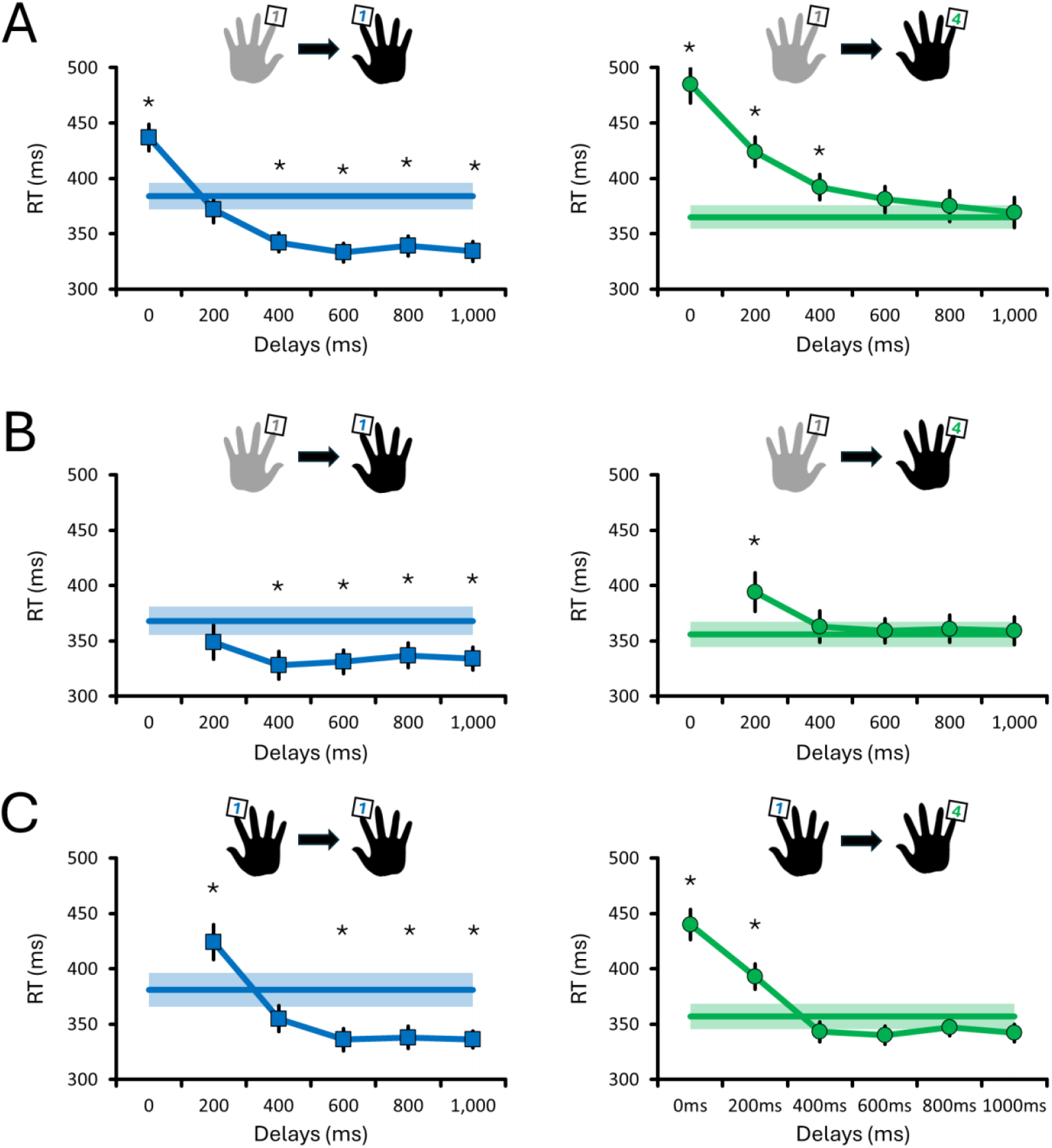
Reaction times (RTs) for right-hand responses following a priming press at varying delays (0–1,000ms). Panels show RTs for homologous (blue squares) and non-homologous actions (green circles) in **(A)** bimanual (Exp 1), **(B)** bimanual without simultaneous presses (Exp 2), and **(C)** unimanual (Exp 3) conditions. Model-derived estimated marginal means ± SE are shown. The horizontal bold lines with shaded errors represent the baseline RTs from unimanual presses. Asterisks (*) indicate significant differences from baseline.

RTs for non-homologous actions were also delay-dependent (χ^2^ = 127, df = 6, *p* < 0.001), but no RT facilitation was observed. Instead, the right little finger showed RT slowing of ~120ms to 30ms for delays ≤400ms (all *p* < 0.034), returning to baseline by ≥600ms (all *p* > 0.101). This indicates that a priming action can interfere with – but not facilitate – the preparation of a subsequent non-homologous response.

### Exp 2: Did interference and facilitation persist when simultaneous bimanual actions are removed?

Exp 2 removed simultaneous presses (0ms delay) to test whether interference and facilitation require synchronous recruitment of both hemispheres. As shown in **Figure 2B**, RTs for homologous presses again varied with the delay between actions (χ^2^ = 82, df = 5, *p* < 0.001). RTs of the right index finger remained near baseline at 200ms (*p* = 0.059) but were ~35ms faster for delays ≥400ms (all *p* < 0.001). This demonstrates that RT facilitation between homologous actions persists even when simultaneous actions are not required.

As in Exp 1, non-homologous presses revealed patterns of RT interference (χ^2^ = 43, df = 5, *p* < 0.001). RTs of the right little finger slowed by ~40ms at 200ms (*p* = 0.005) but returned to baseline for delays ≥400ms (all *p* > 0.761). This shows that interference between non-homologous actions can still emerge in the absence of simultaneous bimanual actions.

### Exp 3: Can interference and facilitation emerge when using a single effector?

Exp 3 eliminated bimanual coordination altogether to test whether RT facilitation occurs when using a single hand. **Figure 2C** shows that even when using a single effector (right index finger), RTs for homologous presses varied with delay (χ^2^ = 72, df = 5, *p* < 0.001). RTs slowed by ~40ms at 200ms (*p* = 0.003), returned to baseline by 400ms (*p* = 0.057), and became ~45ms faster than baseline for delays ≥600ms (all *p* < 0.004). This shows that sequential homologous actions within the same hand can produce RT facilitation.

RTs for non-homologous presses also varied with delay (χ^2^ = 215, df = 6, *p* < 0.001). Simultaneous presses slowed RTs by ~80ms, while 200ms delays resulted in ~40ms slowing (both *p* < 0.001). RTs returned to baseline for delays ≥400ms (all *p* > 0.146). This suggests that even within a single effector, short delays between non-homologous presses produce interference, but not facilitation.

### Control analyses: Was the right little finger responding as fast as the right index finger?

Baseline RTs during unimanual presses showed that the right index (379ms) and little fingers (361ms) responded at comparable speeds across all three experiments (all *p* ≥ 0.060; see **Supplementary Results** for more details). These values represent grand averages across experiments, with each experiment’s mean ± SE shown as horizontal bold lines with shaded error bars in **Figure 2**. Collectively, these baseline RTs indicate that inter-finger differences cannot explain the interference and facilitation effects.

### Control analyses: Did delays also affect RTs of the priming presses?

As a control analysis, we examined RTs of the priming presses across delays (0, 200, 400, 600, 800, and 1,000ms). These results revealed that these RTs were systematically slower whenever a subsequent (or concomitant) press was required, compared with baseline unimanual presses (**Supplementary Results and Figure 1**). This was evident whether priming presses were performed with the left (Exps 1 and 2) or the right index finger (Exp 3). This shows that planning for a second additional action systematically slows the first one, even though the effector and required response remained constant across conditions.

### Control analyses: Did the RT facilitation depend on the actual delays between presses?

RT facilitation was initially analysed using the interval between “Go” stimuli. However, because actual press timings (RTs) varied across trials, the effective delay between actions – the inter-press interval (IPI) – could differ from the scheduled delay between Go cues (**Supplementary Table 1**). This raised the question: is facilitation better explained by the Go stimulus interval or by the IPI? To clarify this, we reanalysed right-hand RTs by grouping trials according to the IPI between the priming and subsequent right-handed presses (**Supplementary Table 2**).

**Supplementary Figure 2** reveals a pattern closely mirroring the main results: regrouping trials based on the IPI did not abolish the facilitation pattern between homologous actions. For non-homologous actions, interference remained pronounced at short IPIs and gradually diminished at longer intervals. These findings confirm that facilitation and interference reflect systematic effects of the temporal delays between actions, rather than artefacts of variability in response timing.

## DISCUSSION

Here, we show that behavioural interference and facilitation during bimanual actions depend on both effector homology and the temporal delay between them. We assessed these effects by indexing RT changes relative to baseline unimanual presses, with slowing interpreted as interference and speeding as facilitation.

For homologous actions, interference emerged at short delays (≤200ms) but transitioned into facilitation at longer delays (≥400ms). Crucially, this facilitation did not require simultaneous engagement of both hands and was also observed when homologous actions were performed with the same hand. Therefore, we propose that this time-dependent facilitation does not strictly rely on interhemispheric interactions.

For non-homologous actions, interference dominated at delays ≤400ms. This effect was observed regardless of whether actions were performed bimanually or unilaterally. Notably, facilitation never occurred for non-homologous actions, suggesting it is specific to homologous responses.

### Time-dependent facilitation of homologous actions

A key novel finding is that facilitation between homologous actions emerged consistently at delays ≥400ms. Notably, facilitation appeared as early as 200ms when RTs were grouped based on the inter-press interval rather than the timing of visual Go stimuli (**see Supplementary Results**). Three observations indicate that this facilitation reflects motor-related dynamics rather than perceptual mechanisms. First, facilitation persisted even when analyses controlled for the effective delays between presses (i.e., inter-press intervals), suggesting that perceptual processes do not fully explain this effect [25,26]. Second, facilitation was absent for non-homologous presses despite identical perceptual demands to homologous ones, further emphasising that facilitation is caused by effector homology and not by perceptual mechanisms. Third, facilitation occurred when processing two Go stimuli compared with the unimanual condition, in which only one stimulus was processed. This pattern indicates that facilitation arose despite the additional demands of processing two stimuli or managing dual-task cognitive load. Interestingly, previous work proposed that interhemispheric facilitation [7–9] underlies enhanced force coupling – whereby force production in one limb enhances output in the other – during simultaneous bimanual activation of homologous effectors [11,12]. Here, we extend this evidence by showing that facilitation persists despite temporal delays, suggesting a functional role for interhemispheric facilitation that extends beyond simultaneous force production.

What mechanism might explain this facilitation? Facilitation may originate from subcortical [19,27] or spinal [28] networks, but cortical signatures of action preparation provide a compelling explanation [16–18]. Electroencephalography studies showed that beta-band power remains suppressed in bilateral sensorimotor areas for a few seconds after a unimanual index finger press [16,17], and until the second action of a sequence has been executed [18]. Since this bilateral beta suppression reflects a release from GABA_A_-mediated inhibition [29,30], one possibility is that a first action disinhibits homologous effectors bilaterally even in the absence of overt bimanual actions [10]. We hypothesise that this disinhibition is specific to homologous actions due to strong transcallosal coherence [31] and recurrent intra-hemispheric projections [15] linking homologous representations within and across hemispheres. The weaker connectivity between non-homologous effectors may explain why they do not benefit from the same disinhibitory dynamics.

The timing and nature of this facilitation remain intriguing. If bimanual disinhibition underlies the effect, why would it initially manifest as interference (≤200ms) before transitioning into facilitation (≥400ms)? Moreover, what exactly is being facilitated: effector identity [13,14], movement directional tuning [32], and/or movement intentions [33]? Homotopic projections are most dense between premotor areas [13,14], suggesting that homologous facilitation occurs during action preparation rather than execution [10]. Moreover, our primarily recruited intrinsic hand muscles; whether facilitation would extend to proximal muscles of the upper limb (e.g., flexor carpi radialis, biceps brachii, deltoid, etc.) remains unclear (see ref [34]). Future research should clarify the anatomical and functional origins of this facilitation.

### Interference – but no facilitation – for non-homologous actions

Another novel finding is that non-homologous actions consistently produced interference at short delays (≤400ms), consistent with previous behavioural studies [2–5]. The interference at short delays is reminiscent of the psychological refractory period [4,5], in which processing a second stimulus is postponed until central resources become available. Here, the interference persisted even when simultaneous bimanual actions were removed and when a single effector was used, suggesting that coordinating non-homologous actions incurs a cost independent of the effector engaged (**Figure 2C**). In contrast to previous models [1,6], we propose that interhemispheric competition alone cannot fully explain this interference. Instead, it may reflect a general central bottleneck [4,5] that taxes neural resources irrespective of whether actions are distributed across hemispheres or confined to a single one. Consistent with this interpretation, our control analyses revealed that even the priming presses were systematically slowed whenever a subsequent response was required (**Supplementary Figure 1**). This suggests that interference is not restricted to the second action, but reflects a broader dual-action cost that influences the initiation of both responses. Importantly, however, bottleneck accounts predict only interference and recovery to baseline, not the facilitation we observed for homologous actions. Future research could examine whether this interference and bottleneck are structural [13,14] and/or functional [4,5], and whether they can be overcome through training [35].

### Beyond interhemispheric inhibition: A shift in the framework

Our findings challenge the long-standing view that bimanual coordination is governed primarily by interhemispheric competition [1,6], and extend classical models that emphasise a preference for spatial and temporal symmetry [11,19,20]. While these models account for competitive neural dynamics and behavioural coupling via symmetry, they do not predict the emergence of time-dependent facilitation when homologous actions are executed sequentially. Our results advance these frameworks on two fronts.

First, interference is not inevitable: it appears at short delays but transitions to facilitation at longer ones – a dynamic, time-sensitive effect not accounted for in current models. Second, facilitation arose in both bimanual and unimanual contexts, indicating it does not strictly depend on interhemispheric pathways. Although this directly challenges the necessity of interhemispheric coupling, we cannot exclude that subcortical structures such as the basal ganglia or cerebellum mediate such interhemispheric-independent homologous facilitation [19,27]. Still, it appears that effector homology shapes sequential actions in ways that have gone largely unrecognised, until now.

These findings carry implications for clinical populations with unilateral motor deficits, such as motor neglect, post-stroke hemiparesis, or apraxia. Although speculative, they raise testable predictions for translational contexts: initiating a homologous action 400 to 1,000ms prior to the impaired response may help recruit compromised or underperforming lateralised motor networks. Before clinical applications can be considered, however, it remains to be determined whether homologous facilitation extends to more complex actions involving the upper and lower limbs.

## CONCLUSION

In sum, this study shows that action homology and temporal delays allow the motor system to flexibly transition between interference and facilitation during bimanual actions. These findings not only refine our understanding of bimanual motor coordination but also open new avenues for enhancing motor performance and rehabilitation strategies through targeted manipulation of timing and effector similarity.

## Supporting information

Supplementary Results

