## Supplementary Results for "Time-Dependent Facilitation of Homologous Actions"

### The right little finger responded as fast as the right index finger

We assessed whether there were differences in the RTs of the right little and right index fingers on unimanual trials. Overall, the results revealed that RTs tended to be 10 to 25ms faster for the right little finger compared with the right index finger, although this difference never reached significance:

- In Exp 1, RTs of the right little finger (365 ± 11ms) did not significantly differ from those of the right index finger (386 ± 12ms; $\chi^2 = 3.5$, df = 1, $p = 0.061$).
- In Exp 2, RTs of the right little finger (357 ± 12 ms) were not statistically faster than those of the right index finger (368 ± 13 ms; $\chi^2 = 1$, df = 1, $p = 0.303$).
- Similar to Exp 1, RTs from Exp 3 also revealed that the right little finger (357 ± 12 ms) was not significantly faster than the right index finger (382 ± 16 ms; $\chi^2 = 3.5$, df = 1, $p = 0.060$).

Altogether, these results indicate that baseline RTs did not significantly differ between the right little and right index fingers, suggesting that the interference and facilitation effects described in the main manuscript cannot be explained by inter-finger differences.

### Left RTs were systematically slower on bimanual *versus* unimanual trials

These analyses assessed whether delays affected simple RTs from the priming finger presses. As in the main analyses, the RTs from the unimanual priming presses – without a concomitant or subsequent press – were used as the baseline. Since priming presses were invariant across trials, regardless of whether a homologous or non-homologous press followed, homology was not included as a fixed factor in the analyses. Results are shown in **Supplementary Figure 1**.

Manipulating delays invariably slowed RTs of the priming left index presses for both Exp 1 ($\chi^2 = 370$, df = 6, $p < 0.001$) and Exp 2 ($\chi^2 = 28$, df = 5, $p < 0.001$). In both cases, RTs were systematically slower on bimanual trials than on unimanual trials, irrespective of the delay (all $p < 0.001$; **Supplementary Figure 1A-1B**). For Exp 3, performing two finger presses with the right hand also slowed RTs from the priming presses ($\chi^2 = 130$, df = 6, all $p < 0.001$; **Supplementary Figure 1C**) at every delay (all $p < 0.001$). Overall, priming presses consistently exhibited longer RTs when a subsequent – or concomitant – press had to be performed.

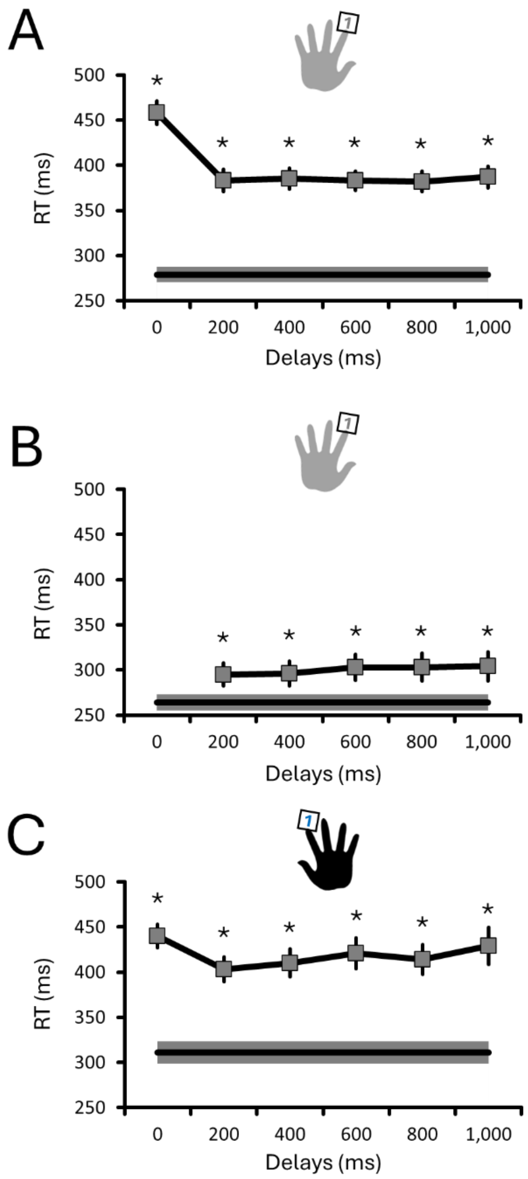


**Supplementary Figure 1. RTs for priming responses at varying delays (0-1,000ms).** Panels show RTs for priming presses in **(A)** bimanual (Exp 1), **(B)** bimanual without simultaneous presses (Exp 2), and **(C)** unimanual (Exp 3) conditions. Model-derived estimated marginal means ± SE are shown. The horizontal bold lines with shaded errors represent the baseline RTs from unimanual presses. Asterisks (*) indicate significant differences from baseline.

### Reanalysing data based on the IPI did not abolish RT facilitation

To assess whether RT facilitation depends on the actual (observed) delay between finger presses – the inter-press interval (IPI) – rather than the interval between "Go!" stimuli, we reanalysed the data by binning trials according to their IPI (**Supplementary Table 1**). Specifically, IPIs between 101–300ms were binned as 200ms delays, IPIs between 301–500ms as 400ms delays, and so forth. **Supplementary Table 2** reports the number of trials that were regrouped per condition after binning according to the above IPI ranges. Supplementary Table 2 reports the mean (± SE) IPI (ms) for each condition of each experiment. Overall, this reanalysis produced a pattern of results nearly identical to the original "Go stimuli"-based analyses (**Supplementary Figure 2**).

For homologous presses, delays affected RTs in all experiments (Exp 1; $\chi^2 = 178$, df = 6, $p < 0.001$ – Exp 2; $\chi^2 = 30$, df = 5, $p < 0.001$ – Exp 3; $\chi^2 = 32$, df = 5, $p < 0.001$). For all experiments, RT facilitation was systematically observed for IPIs ≥400ms (all $p < 0.007$). In bimanual conditions (Exp 1 and Exp 2; **Supplementary Figure 2A and 2B)**, RTs facilitation now emerged as early as IPIs of 200ms (both $p < 0.001$). This earlier onset of facilitation was absent in the unilateral condition (Exp 3; $p = 0.128$; **Supplementary Figure 2C**). This suggests that RT facilitation reflects motor-execution dynamics, as it appears to depend on the effective temporal delay (the IPI) between actions.

For non-homologous presses, delays also affected RTs in all experiments (Exp 1; $\chi^2 = 178$, df = 6, $p < 0.001$ – Exp 2; $\chi^2 = 27$, df = 5, $p < 0.001$ – Exp 3; $\chi^2 = 130$, df = 6, $p < 0.001$), which globally revealed RT interference at short IPIs. In Exp 1 (**Supplementary Figure 2A**), interference was observed for IPIs ≤400ms (all $p < 0.026$). This effect was no longer significant when simultaneous presses were removed (all $p > 0.060$; Exp 2; **Supplementary Figure 2B**) and was only present at 0ms in the single-effector condition ($p < 0.001$; Exp 3; **Supplementary Figure 2C**). These results reinforce that RT interference is predominant when performing actions in close temporal proximity, particularly when actions are executed by separate effectors.

### Supplementary Table 1 - Inter-Press Interval per Condition (ms)

| | | Unimanual | 0ms | 200ms | 400ms | 600ms | 800ms | 1,000ms |
|---|---|---|---|---|---|---|---|---|
| Exp 1 | Homologous | N/A | 13 (1) | 190 (6) | 357 (8) | 553 (10) | 758 (10) | 947 (10) |
| | Non-Homologous | N/A | 14 (1) | 245 (7) | 411 (9) | 602 (9) | 796 (11) | 984 (10) |
| Exp 2 | Homologous | N/A | N/A | 260 (9) | 435 (11) | 626 (10) | 832 (11) | 1029 (15) |
| | Non-Homologous | N/A | N/A | 304 (9) | 471 (8) | 660 (9) | 860 (11) | 1056 (12) |
| Exp 3 | Homologous | N/A | N/A | 224 (4) | 344 (9) | 511 (13) | 714 (16) | 886 (20) |
| | Non-Homologous | N/A | 14 (1) | 193 (7) | 337 (11) | 522 (13) | 722 (15) | 897 (20) |

*The descriptive statistics represent the mean (SE) inter-press interval in ms.*

### Supplementary Table 2 - Change in the Number of Trials per Condition When Regrouping per IPI - Right-Handed Presses

| | | Unimanual | 0ms | 200ms | 400ms | 600ms | 800ms | 1,000ms |
|---|---|---|---|---|---|---|---|---|
| Exp 1 | Homologous | N/A | 0.0 (0.0) | 7.3 (1.6) | 1.2 (0.9) | 0.5 (0.9) | 0.3 (1.0) | -9.3 (1.6) |
| | Non-Homologous | N/A | 0.0 (0.0) | -3.5 (1.8) | 3.7 (1.3) | 1.4 (1.1) | 2.1 (0.8) | -3.6 (1.6) |
| Exp 2 | Homologous | N/A | N/A | -5.6 (2.6) | 1.0 (1.1) | 0.3 (0.9) | 0.4 (1.4) | 4.0 (1.7) |
| | Non-Homologous | N/A | N/A | -14.8 (2.5) | 5.4 (1.1) | -0.7 (1.8) | 0.5 (0.9) | 9.6 (1.9) |
| Exp 3 | Homologous | N/A | N/A | 12.0 (2.6) | 5.2 (1.3) | -1.2 (1.1) | -3.7 (1.2) | -12.3 (2.1) |
| | Non-Homologous | N/A | 0.0 (0.0) | 13.5 (3.0) | 3.0 (0.8) | -2.2 (0.9) | -1.6 (1.1) | -13.1 (1.7) |

*The descriptive statistics represent the mean (SE) change in the number of valid trials per condition for right-handed presses only. Positive and negative numbers reflect an increase and decrease in the number of trials per condition when regrouping per IPI, respectively.*

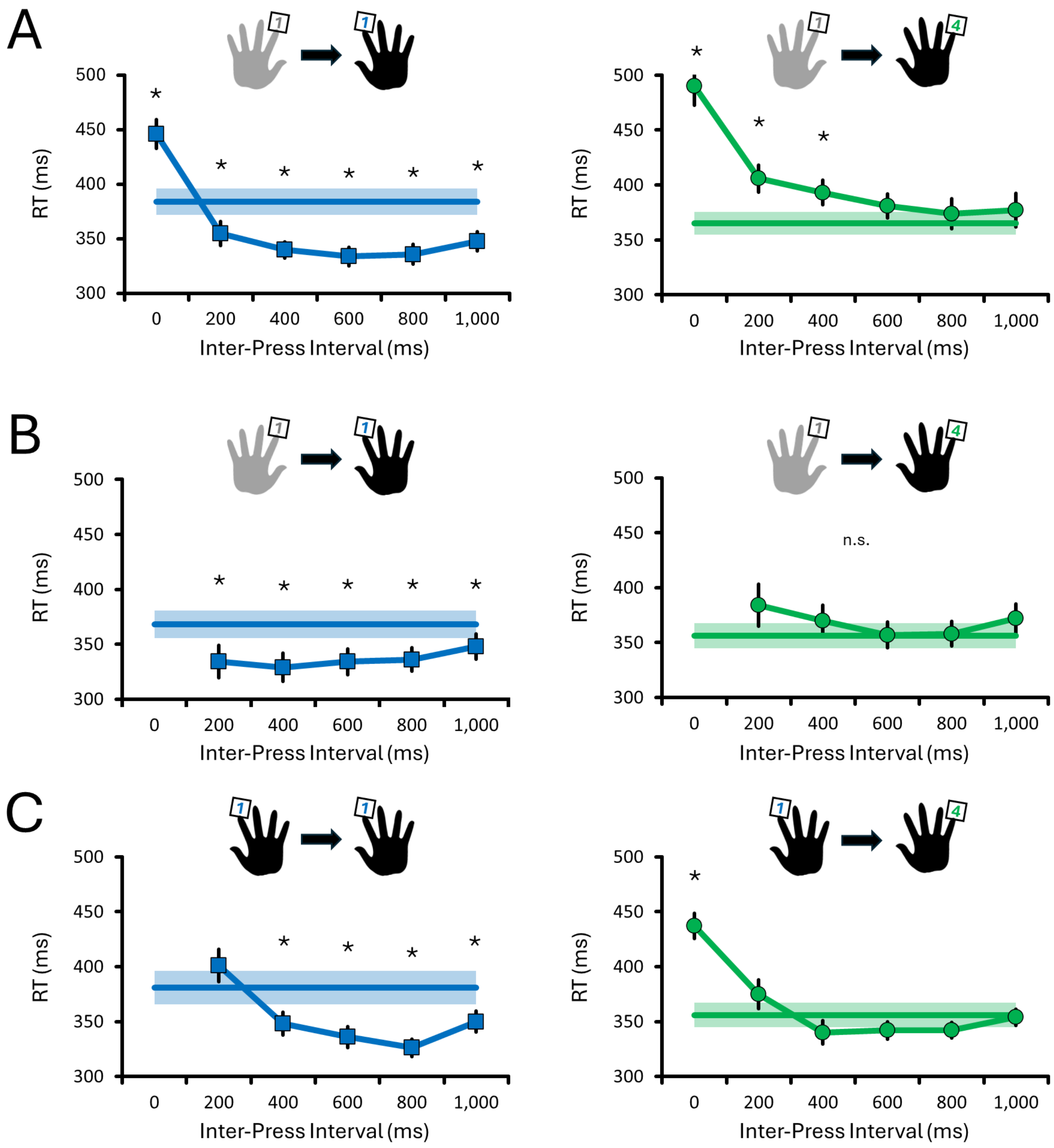


**Supplementary Figure 2. RTs for right index and little finger responses, grouped by inter-press intervals (IPIs).** For homologous actions, RT facilitation emerged at ≥200ms in bimanual conditions (Exps 1 and 2; **A and B**), and ≥400ms in the single-effector condition (Exp 3; **C**). For non-homologous actions, RT interference was observed at delays ≤600ms during bimanual conditions **(A)** but was reduced when simultaneous presses were removed **(B)** or when actions involved a single effector **(C)**. Model-derived estimated marginal means ± SE are shown. The horizontal bold lines with shaded errors represent the baseline RTs from unimanual presses. Asterisks (*) indicate significant differences from baseline.
